# Krüppel-like Factors Play Essential Roles in Regulating Pluripotency and the Formation of Neural Crest Stem Cells

**DOI:** 10.1101/2025.01.13.632647

**Authors:** Sara Rigney, Joshua R. York, Carole LaBonne

## Abstract

The evolutionary transition from simple chordate body plans to complex vertebrate body plans was driven by the acquisition of the neural crest, a stem cell population that retains broad, multi-germ layer developmental potential long after most embryonic cells have become lineage restricted. We have previously shown that neural crest cells share significant gene regulatory architecture with pluripotent blastula stem cells. Here we examine the roles that Krüppel-like Family (Klf) transcription factors play in these stem cell populations. Although Klf4 has established roles in regulating pluripotency in mammalian stem cells cultures, we find that in *Xenopus* it is *klf2* that is highly expressed in pluripotent blastula stem cells. *klf2* expression is down-regulated as cells transition to a neural crest state while a related klf factor, klf17, is significantly up regulated in response to neural crest induction. We used gain and loss of function studies to compare the activities of these closely related factors and found that they have both shared and distinct activities. Inhibition of either klf2 or klf17 activity led to significantly expanded expression of pluripotency, neural plate border and neural crest factors in neurula stage embryos, leading us to hypothesize that klf factors regulate the exit from pluripotency and proper establishment of the boundary of the neural crest domain. To gain further insights into the role of klf factors in the evolution of the neural crest, we examined their expression in the jawless vertebrate, *Petromyzon marinus (*sea lamprey). We find that lamprey have a *klf2/4* and a *klf17* gene, but that only *klf17* is expressed in blastula and neural crest stem cells. Moreover, ectopic expression of lamprey *klf17* in *Xenopus* embryos phenocopies *Xenopus klf17* activity. These data suggest that klf17, rather than klf4, may have been the ancestral klf factor that functioned in these GRNs in stem vertebrates.

## Introduction

The neural crest is a stem cell population that is unique to vertebrates that is notable for its broad multi-germ layer developmental potential (Le Douarin and Kalcheim 1999; Prasad, Sauka-Spengler, and LaBonne 2012; Dupin and Le Douarin 2014). Neural crest cells contribute an amazingly diverse array of highly specialized cell types to the vertebrate body plan, including sensory neurons and glia of the peripheral nervous system, pigment-producing melanocytes, and bone and cartilage of the craniofacial skeleton (Groves and LaBonne 2014; Schock, York, and LaBonne 2023). Notably, these cells retain their broad developmental potential even as neighboring cells become lineage restricted. Thus, understanding the developmental origins of neural crest stem cells and their broad potential is key to understanding origin and maintenance of stem cell potential and the early evolution of vertebrates.

New insights into the origins of neural crest potential came from the realization that these cells share significant gene regulatory network (GRN) architecture with the pluripotent cells of vertebrate blastula-stage embryos, including a large cohort of transcription factors (Lavial 2007; Buitrago-Delgado et al. 2015; Lignell et al. 2017; Lukoseviciute et al. 2018; Scerbo 2020; Zalc et al. 2021; Schock, York, and LaBonne 2023). These findings suggest a model where neural crest cells arose via retention of characteristics of those earlier cells. Consistent with such a model, a requirement for BMP signaling, and FGF-mediated MAP kinase signaling have also been found to be shared attributes of pluripotent blastula cells and neural crest cells in *Xenopus* (Geary and LaBonne 2018; Nordin and LaBonne 2014). Both stem cell populations are also characterized by low levels of histone acetylation and a requirement for both histone deacetylase (HDAC) and Brd4 activity, indicating the important roles that epigenetic factors play in these cells (Rao and LaBonne 2018; Huber, Rao, and LaBonne 2024).

Recent comparative genomic work in *Xenopus* and lamprey has demonstrated that a shared pluripotency-neural crest GRN had already been assembled in stem vertebrates (York et al. 2024), providing further evidence that the multi-germ layer potential of the neural crest evolved in early vertebrates by deploying this shared regulatory program. Lampreys, one of only two extant jawless vertebrates (the other being hagfish) have evolved independently from jawed vertebrates for more than 500 million years (Smith et al. 2018). Thus, traits shared between lampreys and jawed vertebrates such as *Xenopus* likely represent those present in the last common ancestor of extant vertebrates. Notably, both the blastula and neural crest GRNs were found to be highly conserved between *Xenopus* and lamprey, even at the level of absolute transcript levels (York et al. 2024). There were, however, a number of notable differences between the lamprey and *Xenopus* GRNs. One of these centered on Krüppel-like factors (Klfs), a large diverse family of transcription factors characterized by three highly conserved zinc fingers in their C-terminal DNA binding domains (Miller 1993; Pearson et al. 2008; Presnell, Schnitzler, and Browne 2015). These factors can act as either transcriptional activators or repressors (Jha et al. 2024). In mammals, Klf2 and Klf4 have established roles in pluripotency in blastula embryos and derived embryonic stem cells (Bieker 2001; Jiang et al. 2008; Bourillot 2010; McConnell and Yang 2010; Yeo et al. 2014; Bialkowska, Yang, and Mallipattu 2017; Lea et al. 2021). Unlike mammals and other amniotes, however, lamprey Klf2/4 is not expressed in blastula or neural crest stem cells (Hockman et al. 2019; York et al. 2024), whereas a closely related factor, *klf17* is expressed..

*Klf4* is one of the original Yamanaka factors, a transcription factor that in combination with *Myc*, *Sox2*, and *Oct4* was shown capable of reverting differentiated somatic cells back to a pluripotent state in mammalian cell culture (Takahashi and Yamanaka 2006). Follow-up studies expanding the network of factors capable of reprogramming somatic cells found that *Klf2* could replace *Klf4* in this role (Nakagawa et al. 2008; Feng et al. 2009; Gillich et al. 2012). Moreover, a Klf regulatory network consisting of *Klf2, Klf4* and *Klf5* is involved in maintaining pluripotency in mouse embryonic stem cells (mESCs) (Jiang et al. 2008; Parisi et al. 2008; Yeo et al. 2014). It has been reported that due to at least partial functional redundancy, knockdown of all three Klf factors is required to induce the differentiation of mESCs (Jiang et al. 2008). Notably, all three Klf factors share key regulatory targets with the pluripotency factor *Nanog*, including *Pou5f1* and *Sox2* (Jiang et al. 2008; Parisi et al. 2008; Bourillot et al. 2009; Hall et al. 2009; Wei et al. 2009; Zhang et al. 2010; Yeo et al. 2014).

Moreover, all three of these Klf factors bind upstream regulatory regions in the *Nanog* promotor and promote its expression (Jiang et al. 2008). Despite these redundancies, genome occupancy experiments indicate that each of these Klf factors also regulate unique targets, and suggest that they function hierarchically to promote pluripotency (Jiang et al. 2008; Parisi et al. 2008; Yeo et al. 2014). Despite their well-studied and critical role in regulating pluripotency in cultured mESCs, the role of Klf factors in regulating pluripotency in blastula stem cells *in vivo,* as well as their role in neural crest cells during development, has remained largely unexplored.

Here we show that of the three klfs required in mESCs, only *klf2* is highly expressed in the blastula stem cells of *Xenopus* embryos. We further show that *Xenopus* blastula stem cells express lower levels of a related factor, *klf17*, which is subsequently expressed at higher levels in the neural plate border and neural crest. We provide evidence that klf2 and klf17 regulate the exit from pluripotency in early embryonic cells and subsequently control the expression boundaries of neural plate border and neural crest genes. Despite the temporal expression differences of *klf2* and *klf17* our functional studies show that the proteins they encode have largely overlapping activities. Finally, we explore the evolutionary origins of klf activity in blastula and neural crest stem cells. We provide evidence for deep conservation of klf17 function in both stem cell populations across cyclostomes and gnathostomes and our data suggest that klf17, rather than *klf2/4,* was likely the ancestral klf factor involved in regulating the developmental potential of blastula and neural crest stem cells in early vertebrates.

## Results

### *klf2* and *klf17* are expressed in blastula and neural crest stem cells

To determine which *klf* family genes are expressed in blastula and/or neural crest stem cells in *Xenopus* embryos we mined our previously published transcriptomes for pluripotent blastula and neural crest stem cells (York et al. 2024) for all known klf factors in the *Xenopus* genome. In contrast to mESCs (Jiang et al. 2008; Parisi et al. 2008; Yeo et al. 2014) we found that *Xenopus klf2* was the most highly expressed klf factor in blastula animal pole cells (Fig. S1A), and its expression was approximately 17 fold higher than *klf4*. When we examined the expression *klf* factors as blastula cells were induced to adopt a neural crest state we found that *klf17* was the only family member whose expression increased in early (stage 13) neural crest cells. Additionally, it was the most highly expressed *klf* factor in both stage 13 and stage 17 neural crest cells (Fig. S1A). By contrast, *klf2* expression was reduced in neural crest cells relative to its blastula expression (Fig. 1A). While *klf17* was also expressed in blastula stem cells, its expression was significantly (2.4 fold; p =0.002) lower level than that of *klf2*.

**Fig 1.**
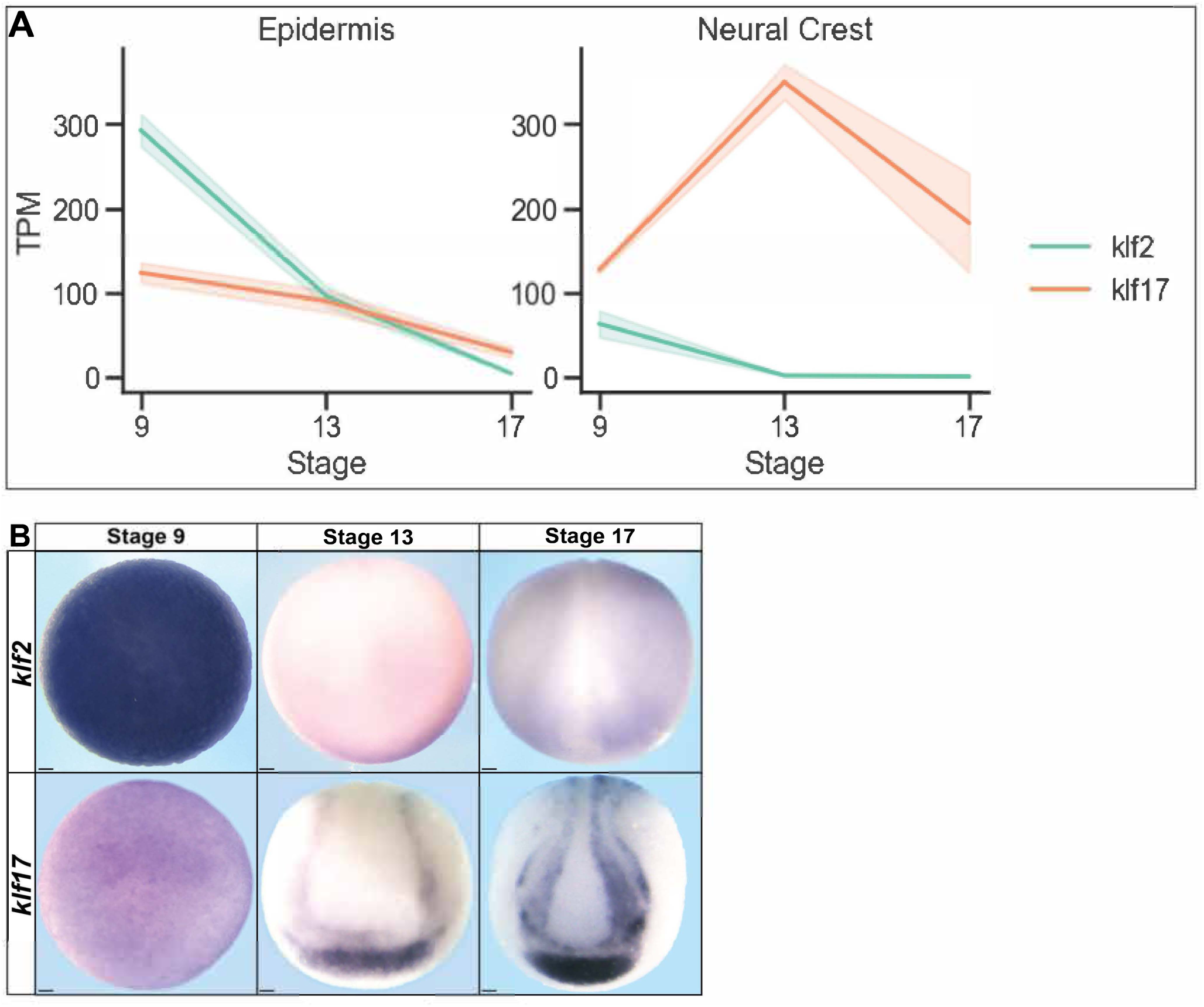
*klf2* and *klf17* are dynamically expressed in the blastula, neural plate border and neural crest cells of xenopus explants and embryos. (A) Graph depicting average TPMs of *klf2* and *klf17* in epidermis and neural crest explants at stages equivalent to blastula (stage 9), early neurula (stage 13), and late neurula (stage 17) of whole embryos. Error bars represent the standard error of mean. (B) *In situ* hybridization examining *klf2* and *klf17* expression in whole mount wildtype Xenopus embryos collected at blastula (stage 9), early neurula (stage 13) and late neurula (stage 17) stages. Scale bars: 100 µM.

We next examined the temporo-spatial expression of *klf2* and *klf17* in *Xenopus* embryos using whole mount in situ hybridization (WISH). Consistent with our transcriptomic data we observed high expression of *klf2* in blastula animal pole cells and significantly lower expression of *klf17*. At early neurula stages (stage 13) *klf17* is expressed at the neural plate border and the presumptive cement gland. *klf2* expression is undetectable by WISH at state 13, although transcripts are detected by RNA-Seq. By stage 17 low levels of *klf2* expression are observed throughout the non-neural ectoderm. Interestingly, by this stage, *klf17* expression in the neural folds has resolved to distinct medial and lateral domains (Fig. 1B).

We next used hybridization chain reaction (HCR) probes to further examine the expression of *klf17* at neurula stages and compare its expression to canonical neural plate border and neural crest factors *pax3* and *snai2*, respectively. We observed significant overlap of *klf17* and *pax3* expression at the neural plate border at stage 13 (Fig. 2 A,B). While *snai2* expression is much lower at this stage, most *snai2-*positive cells also expressed *klf17* (Fig. 2 C,D). Strikingly, however, by stage 17 the medial and lateral *klf17* expression domains surrounded the *snai2* domain but did not overlap with it. Cells within the medial band of Klf17 expression still co-expressed *pax3* at this stage (Fig. 2 A,B). We used line profile averages of confocal images to further illustrate these changes that occur to the spatial relationships between *klf17* and *pax3*, and *klf17* and *snai2* expression domains during the establishment of the neural plate border and neural crest (Fig. 2 B,D)

**Fig 2.**
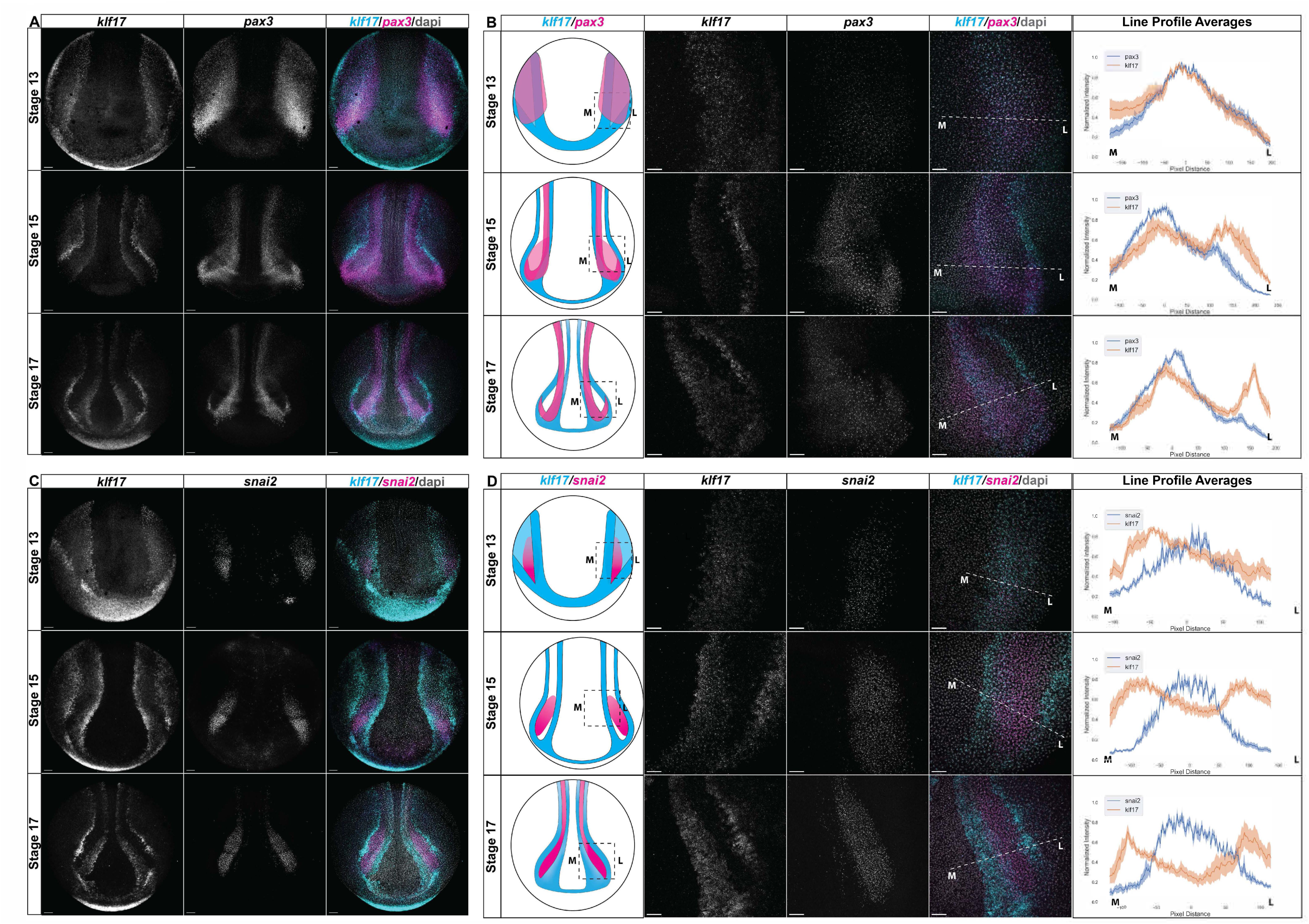
Overlapping expression patterns of *klf17* with neural plate border and neural crest changes during neurulation. (A,C) Whole embryos probed with HCR oligos examining the expression patterns of (A) *klf17* (cyan) and the neural plate border marker *pax3* (magenta) or (C) *klf17* (cyan) and the neural crest marker *snai2* (magenta) at early (stage 13), mid (stage 15), and late (stage 17) neurulation. (B,D) Average line profile measurements of the normalized pixel intensity values across the medial-lateral axis of maximum projection images of the neural plate border and neural crest cell region of wild type embryos. Diagrams in the first column represent the embryos and gene expression patterns at each neurula stage analyzed. The dashed box on the diagram depicts the zoomed in area of the embryo imaged for each line profile measurement. Representative individual (B) *klf17* and *pax3* or (D) *klf17* and *snai2* HCR images are shown respectively in columns two and three as well as the merged image in column four. The dotted line on the merged image depicts the axis the line profile averages were measured across with “M” being the medial end point and “L” the lateral endpoint in respect to the embryo midline. The final column plots the average line profile measurements for each stage (minimum n=5). The x-axis plots the pixel positions along the measurement axis, with the peak of either *pax3* (B) or *snail2* (D) profiles centered at 0 (after curve fitting). The y-axis plots the average normalized intensity values of either (B) *pax3* (blue) and *klf17* (orange) or (D) *snai2* (blue) and *klf17* (orange) with shading measuring the standard error of mean. Scale bars: 150 µM..

### klf2 or klf17 depletion expands the plate border and neural crest domains

Having characterized the expression of *klf2* and *klf17* we next asked if these factors are required for establishment of the neural plate border and/or neural crest. To this end, we injected 2 cells of 8-cell embryos with translation-blocking morpholinos (MO) specific to each of these klf factors (Fig. S2A) and cultured them to neurula stages for WISH. We found that MO-mediated depletion of either klf2 or klf17 resulted in expanded expression of the neural plate border markers zic1 (klf2MO: 90.5%, n=74; klf17MO 88.3%, n=77), pax3 (klf2MO: 86.2%, n=65; klf17Mo: 83.1%, n=83), msx1(klf2MO: 75%, n=60; klf17MO: 85%, n=60) relative to the uninjected side (Fig. 3A). As neural plate boarder factors are required for formation of definitive neural crest cells (Luo et al. 2002; Luo 2003; Monsoro-Burq, Wang, and Harland 2005; Groves and LaBonne 2014; Simões-Costa and Bronner 2015), we next examined the effects of klf2 or klf17 depletion on the expression of neural crest factors. Depletion of either klf2 or klf17 resulted in modest expansion of the expression domains of *snai2* (klf2MO: 77.1%, n=166; klf17MO: 68.4% n=177) and *foxd3* (klf2MO: 70.8%, n=106; klf17MO 73.5%, n=136) (Fig. 3B) that was less pronounced than that observed for neural plate border factors. Together, these findings indicate that despite the temporal differences in their expression, both *klf2* and *klf17* are required for the proper establishment of the spatial boundaries of both the neural plate border and neural crest.

**Fig 3.**
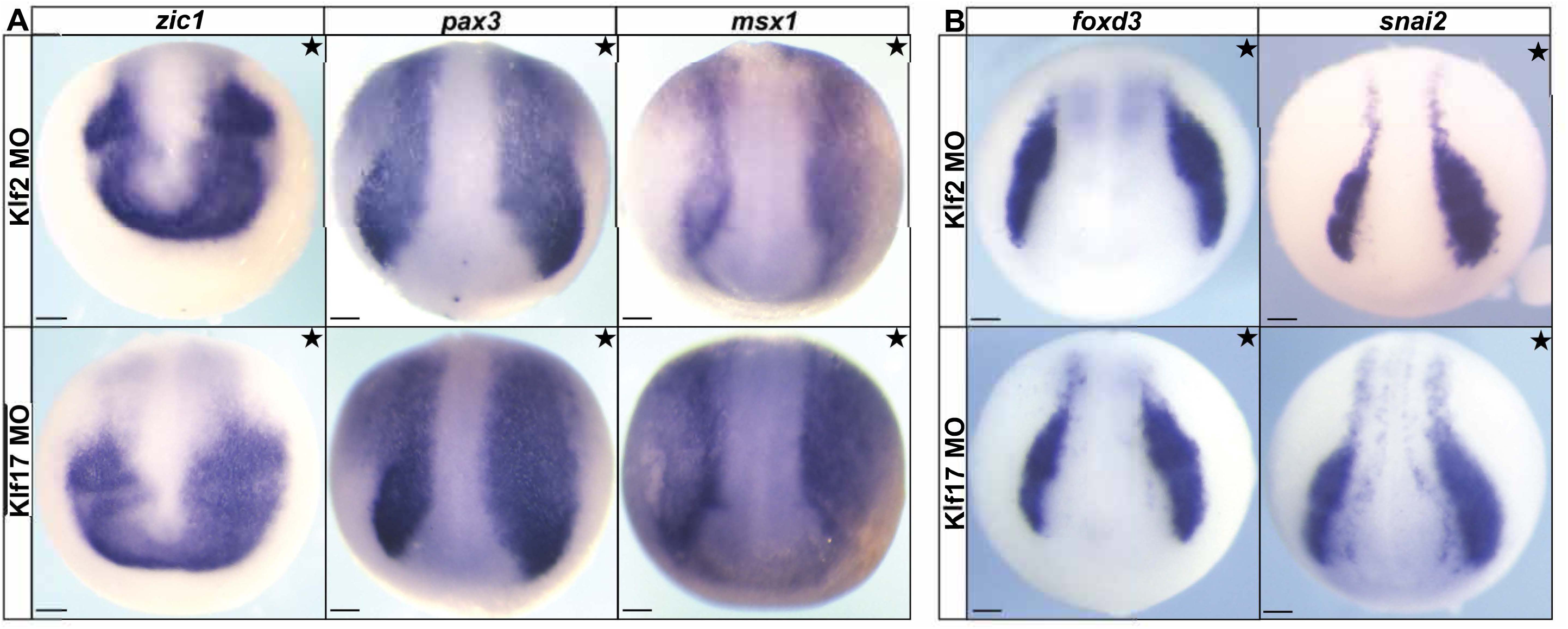
*klf2* and *klf17* are essential to establish the proper neural plate border and neural crest domain boundaries. **(A,B)** *In situ* hybridizations of embryos unilaterally injected with either *klf2* or *klf17* fluorescein-tagged morpholinos (stars denotes the injected side). The fluorescein tag was used as a lineage tracer and embryos were pre-sorted for left/right targeting. (A) Neural plate border markers *pax3*, *zic1*, and *msx1* in early neurula embryos. (B) Neural crest cell markers *foxd3* and *snai2* in late neurula embryos. MO, morpholino; scale bars: 100 µM.

### *klf2* or *klf17* GOF inhibits expression of neural crest markers

Because the above loss-of-function experiments suggest that klf2 and klf17 act directly or indirectly to restrict the size of neural plate border and neural crest domains, we hypothesized that klf2 and/or klf17 gain-of-function might interfere with expression of neural plate border and/or neural crest factors. To test this, we injected one cell of two-cell stage embryos with mRNA encoding epitope-tagged klf2 or klf17 and cultured embryos to neurula stages for WISH. Consistent with this hypothesis, ectopic activity of either klf2 or klf17 resulted in near-total loss of the neural crest factors *snai2* (klf2nMT: 90.7%, *n*=107; klf17nMT: 84.2%, *n*=114) and *foxd3 (*klf2nMT*: 94.5%, n=91;* klf17nMT*: 93.7%, n=95)* (Fig. 4A). When effects on neural plate border markers *pax3* and *zic1* were examined, the phenotypes were more complex, as both klf factors caused regions of lost expression as well as regions of ectopic expression, primarily in the medial neural plate [*pax3* (klf2nMT: 98.3, *n*=59; klf17nMT: 91.7%, *n*=72) and *zic1* (klf2nMT: 98.7%, *n*=75; klf17nMT: 96.1%, *n*=102)] (Fig. 4B). Together these results suggest that both klf2 and klf17 regulate the spatial boundaries of the neural plate border and neural crest domains.

**Fig 4.**
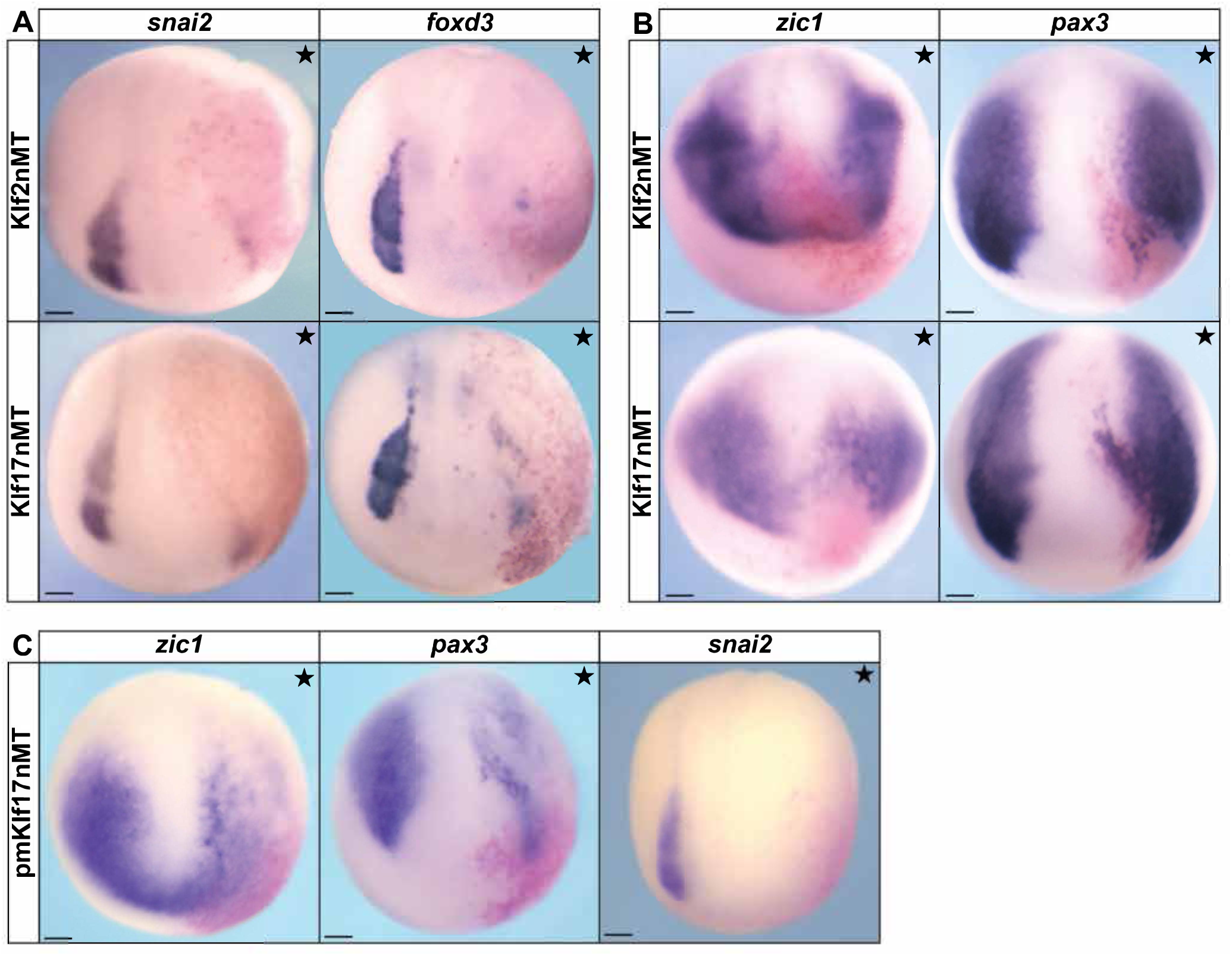
Ectopic effects of klf transcription factors on the establishment of proper neural plate border boundaries and neural crest cell formation are evolutionarily conserved. (A,B) *In situ* hybridizations of embryos unilaterally expressing klf2 or klf17 epitope tagged mRNA (stars denote injected side). β-Galactosidase (red) was used as a lineage tracer. (A) *snai2* and *foxd3* in late neurula embryos. (B) *pax3* and *zic1* in early neurula embryos. (C) *In situ* hybridizations for *zic1, pax3,* and *snai2* in neurula embryos unilaterally expressing *Petromyzon marinus* (sea lamprey) *klf17* epitope tagged mRNA (stars denote injected side). β-Galactosidase (red) was used as a lineage tracer. nMT, n-terminal myc tag; pm, *Petromyzon marinus;* scale bars: 100 µM.

We previously showed that in lamprey, *klf17* is the only *klf2/4/17* clade member expressed in neural crest and blastula stem cells (York et al. 2024)( Fig. S4A). We therefore asked if the ability of klf17 to restrict neural crest might be deeply conserved to the base of vertebrates by performing heterologous gain-of-function experiments. mRNA encoding epitope-tagged lamprey klf17 was expressed unilaterally in two-cell stage *Xenopus* embryos at levels matching those of *Xenopus* klf17. We found that lamprey klf17 largely phenocopied the repressive effects of *Xenopus* klf17, resulting in near-total loss of neural plate border and neural crest gene expression for *pax3 (*100%, n=48), *zic1 (90.2%, n=41)*, and *snai2 (100%, n=61)* (Fig. 4C). These results suggest that ability of klf17 transcription factors to inhibit neural plate border and neural crest has been largely conserved across jawed and jawless vertebrate lineages.

### klf2 and klf17 regulate expression of core pluripotency genes

Pluripotency genes are required for proper establishment of the neural plate border in *Xenopus* as components of the blastula stage pluripotency GRN are coopted for establishment of this domain (Shock et al 2024; York et al 2024). We therefore examined the expression of pluripotency genes during establishment of the neural plate border. We found loss of either klf2 or klf17 resulted in expanded expression of *pou5f3.2* (klf2MO: 64.2%, *n*=53; klf17MO: 81.3%, *n*=64), *sox3* (klf2MO: 93.8%, *n*=48; klf17MO: 91.5%, *n*=47), and *tfap2a* (klf2MO: 95.5%, *n=44;* klf17MO: 83.6%, *n*=55) at these stages (Fig. 5a) suggesting that klf2 and klf17 also play essential roles is determining boundaries of the pluripotency-associated gene expression domains as embryos progress from blastula to neurula stages. This expansion was particularly dramatic for *pou5f3.2* expression, perhaps indicative of a more direct regulation of this target.

**Fig 5.**
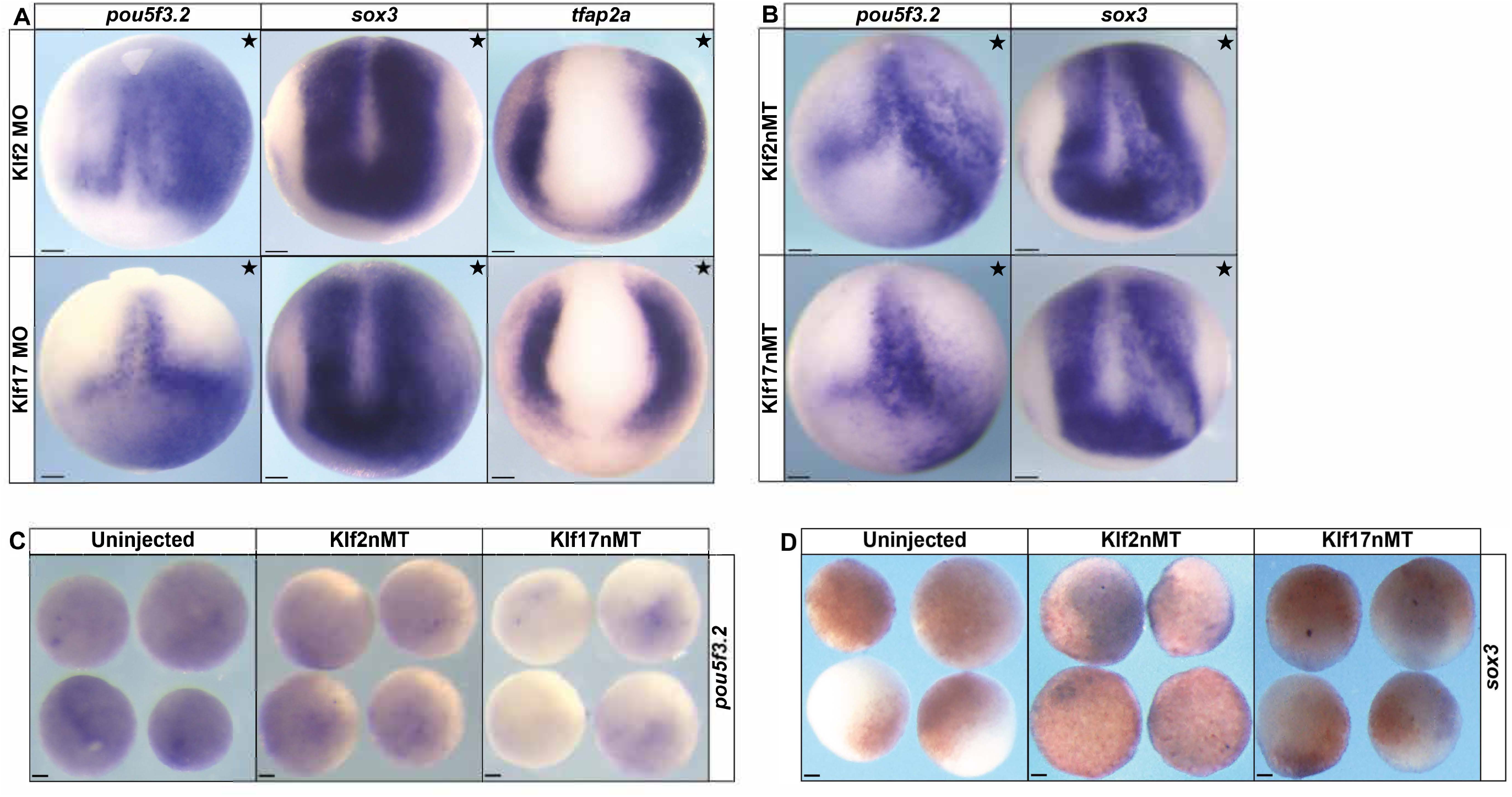
*klf2* and *klf17* regulate pluripotency factors expression and are essential to establish their domain boundaries in neurula-stage embryos. (A) *In situ* hybridizations for *pou5f3.2, sox3,* and *tfap2a* in early neurula embryos unilaterally injected with either *klf2* or *klf17* fluorescein-tagged morpholinos (B) *In situ* hybridizations for *pou5f3.2* and *sox3* in embryos unilaterally expressing klf2 or klf17 epitope tagged mRNA(stars denotes the injected side). The fluorescein tag was used as a lineage tracer and embryos were pre-sorted for left/right side targeting. (C) *pouf5f3.2;* fluorescein dextran was used as a lineage tracer and explants were pre-screened for targeting. (D) *sox3;* β-Galactosidase (red) was used as a lineage tracer. MO, morpholino; nMT, n-terminal myc tag; scale bars: 100 µM.

We also examined the effects of increased klf activity on pluripotency markers (Fig. 5B). While we had expected that klf2 and klf17 would inhibit the expression of markers such as *pou5f3.2* and *sox3*, the actual phenotype was more complex. Similar to their effects on the neural plate border markers *zic1* and *pax3*, both klf2 and klf17 partially inhibit the expression of these factors but also induced regions of ectopic expression (*pou5f3.2*: klf2nMT 91.2%, n=57; klf17nMT 86.5%, n=52; *sox3*: (klf2nMT 87.8%, n=49; klf17nMT 83.3%, n=48). Because components of the pluripotency GRN are known to have complex cross-regulatory interactions, it is likely that klf2/17 gain of function has distinct regional effects depending on what interacting factors are expressed there and at what level.

Given the complexity of the whole embryo phenotype in response to increased klf2 and klf17 activity we turned to explant assays, where the responding tissue would be more uniform, allowing us to more effectively assess the role of Klfs. Two-cell embryos were injected in both cells with mRNA encoding either epitope tagged klf2 or klf17 and cultured to blastula stage when animal pole cells were explanted and further cultured to stage 13 for analysis. No ectopic expression of *pou5f3.2* or *sox3* was observed indicating that the klf-mediated ectopic expression of these factors in whole embryos was indirect (Fig. 5C,D).

### klf2 and klf17 inhibit lineage restriction

In *Xenopus* embryos, pluripotent blastula stems cells can easily be explanted and instructed to give rise to any germ layer. Given no alternative signals, these cells transit to an epidermal stage due to autocrine BMP signaling and will express *krt12.4*. To determine how upregulation of klf2 and klf17 activity would affect the ability of blastula cells to transit toward lineage restriction, two-cell embryos were injected in both cells with mRNA encoding either klf2 or klf17 and cultured to blastula stage when animal pole cells were explanted, then further cultured to stage 13 for analysis. In contrast to control explants, explants expressing either ectopic klf2 or klf17 failed to form epidermis as evidenced by the loss of *krt12.4* (klf2nMT: 81%, n=42; klf17nMT: 87.2%, n=47; WT: 0%, n=34) (Fig. 6A).

**Fig 6.**
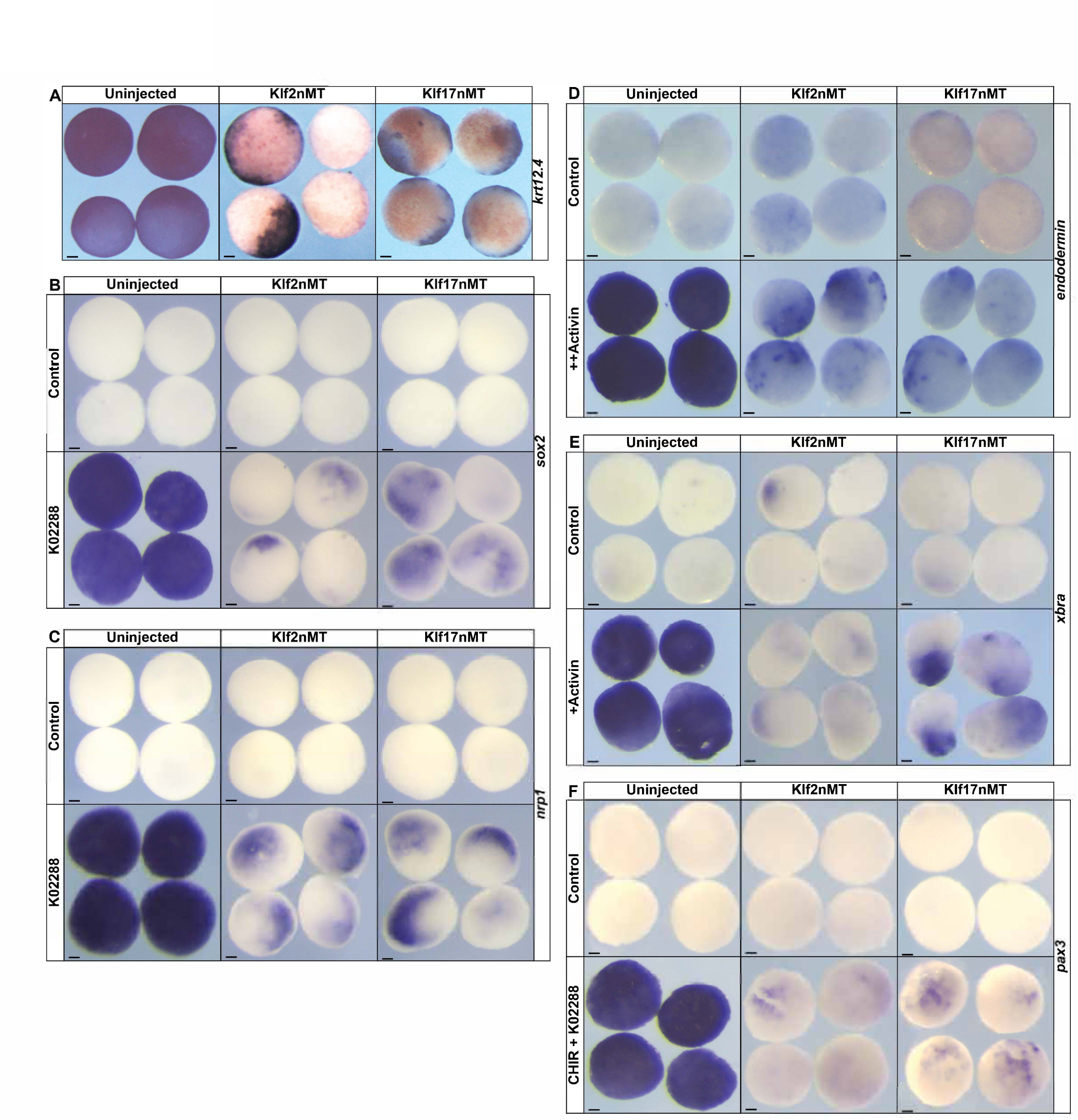
Over expression of klf2 and klf17 impairs lineage restriction. (A-F) *In situ* hybridizations of either uninjected wild type explants or explants expressing *klf2* or *klf17* epitope tagged mRNA examining the expression of: (A) epidermal marker *krt12.4;*(B-C) neural markers *sox2 (B)* and *nrp1 (C*) with or without the BMP antagonist K02288*; (D)* endodermal marker *endodermin* with or without a high dose of activin; (E) mesodermal marker *xbra* with or without a high dose of activin; and (F) neural plate border marker *pax3* with or without a combination of a Wnt agonist (CHIR) and BMP antagonist (K02288) small molecule cocktail. nMT, n-terminal myc tag; K02288, BMP inhibitor; CHIR, CHIR99021 (Wnt agonist); +, low activin; ++, high activin; scale bars: 100 µM.

Treatment with a small molecule antagonist of BMP signaling, K02288, directs blastula explants to adopt a neural fate and express neural progenitor genes such as *sox2* (WT: 100%, *n*=34) and *nrp1* (WT: 100%,*n*=31) (Fig. 6 B,C). Ectopic expression of either klf2 or klf17 prevented explants from adopting a neural fate with an observed loss of *sox2* (klf2nMT: 96.9%, *n*=32; klf17nMT: 90.3%, *n*=31) and *nrp1* (klf2nMT: 100%, *n*=33; Klf17nMT: 93.8%, *n*=32) expression (Fig. 6B,C).

Explants can be induced to form endoderm by treatment with high doses of activin, as evidenced by expression of *endodermin*. Ectopic expression of either klf2 or klf17 blocked *endodermin* induction (klf2nMT: 100%, n=32; klf17nMT: 93.8%, n=32, WT: 0%, n=32) (Fig. 6D). Similarly, treatment with low doses of activin promote mesoderm formation as evidenced by *xbra* expression, and this too was blocked by either klf2 or klf17 (klf2nMT: 90.9%, n=33; klf17nMT: 90%, n=30; WT: 0%, n=34) (Fig. 6E). Finally, we asked if these klf factors would also inhibit neural plate border formation. Explants can be induced to a neural plate border state by treatment with a combination of K02288 and the small molecule wnt agonist, CHIR99021(CHIR) (Huber and LaBonne 2024). Control explants treated with CHIR/K02288 strongly expressed the neural plate border factor *pax3* (WT: 100%, *n*=30) whereas ectopic expression of either klf2 or klf17 resulted in a failure to induce *pax3* expression (klf2nMT: 91.4%, n=35; klf17nMT: 97.2%, n=36) (Fig. 6F). Together these data support a model where up regulation of klf2 or klf17 restricts the pluripotency of blastula explants.

### Loss of klf activity prolongs expression of pluripotency markers expression in blastula explants

The above results demonstrate that klf2 and klf17 negatively regulate expression of neural crest and pluripotency factors and block functional pluripotency. We therefore hypothesized that loss of klf2 and/or klf17 might prolong the ability of pluripotent blastula cells to form different cell types beyond stages when they are normally competent to do so. To test this, 8-cell embryos were injected in all four micromeres with either klf2 or klf17 MOs and animal pole cells were explanted at stage 9 and cultured to stage 13. While control explants had down-regulated expression of *sox3* by this stage, explants depleted for klf2 or klf17 still expressed low levels of *sox3* (klf2MO: 100%, n=34; klf17MO: 100%, n=30; WT: 0%, n=31) (Fig. 7A). We also examined *pou5f3.2* and found that its expression was enhanced in response to either klf2 or klf17 depletion (klf2MO: 93%, n= 43; klf17MO 94.1%, n=34) (Fig. 7B) Consistent with the persistence of *sox3* and *pou5f3.2* expression these explants did not properly lineage restrict, as evidenced by reduced expression of the epidermal marker *krt12.4* (klf2MO: 93.5%, *n*=31; klf17MO: 93.5%, n=31; WT: 0%, n=34)(Fig. 7C). Interestingly Klf depletion did not inhibit CHIR/k02288-mediated reprograming to a neural plate border state (klf2MO: 93.3%, *n*=30; klf17MO: 93.1%, n=29; WT: 100%, n=29) (Fig.7D) likely because of the adjacency of this state to the pluripotent blastula state.

**Fig 7.**
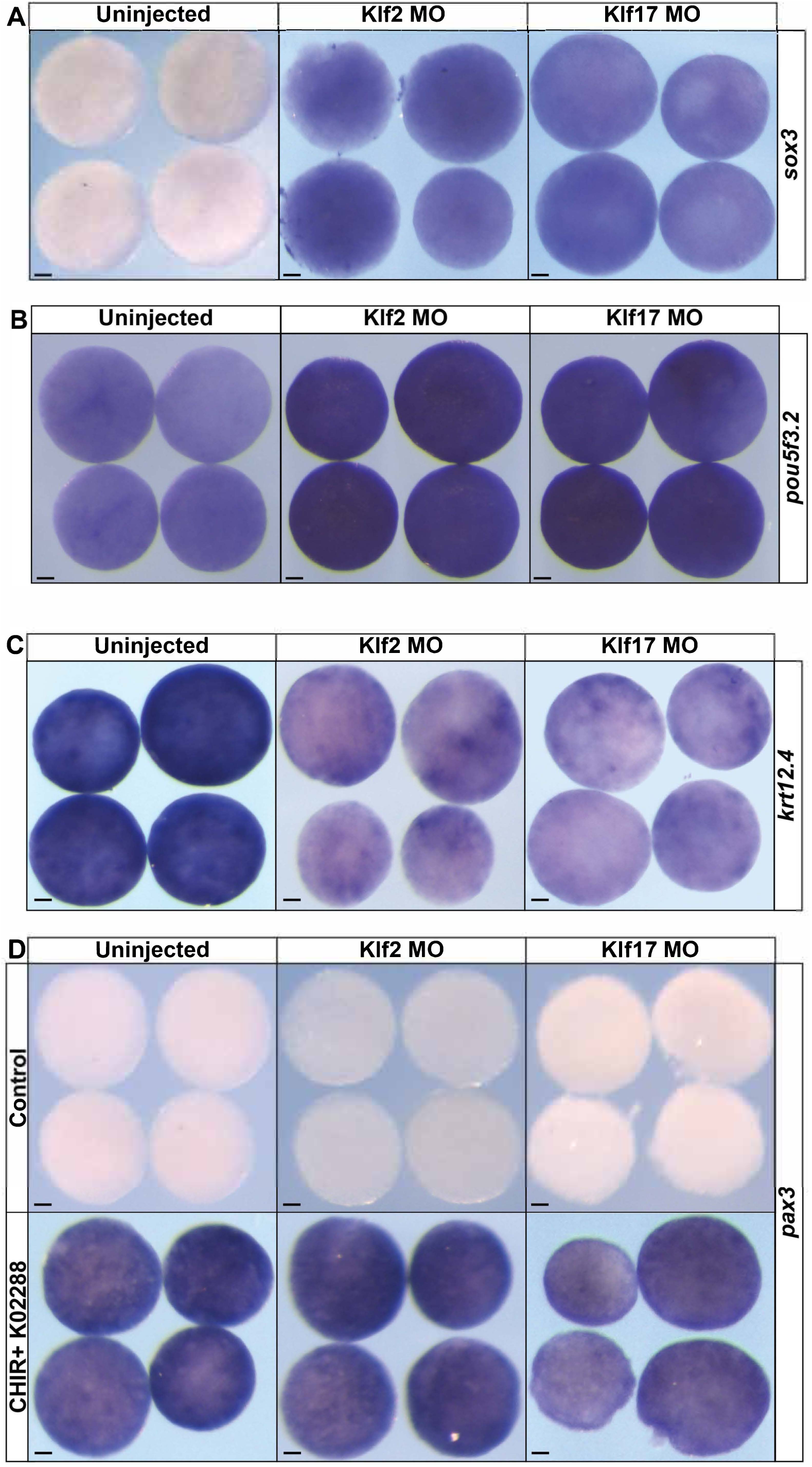
Knockdown of *klf2* or *klf17* prolongs the expression of pluripotency factors. (A-D) *In situ* hybridizations of uninjected wild type, *klf2* morphant, or *klf17* morphant explants (stage 12.5-13) examining the expression of: (A) *sox3;* (B) *pou5f3.2;* (C) *krt12.4;* and (D) *pax3* with or without the CHIR (Wnt agonist) + K02288 (BMP antagonist) small molecule cocktail. MO, morpholino; K02288, BMP inhibitor; CHIR, CHIR99021 (Wnt agonist); scale bars: 100 µM.

### klf2 and klf17 regulate the exit from pluripotency

The expanded expression of pluripotency factors at the neural plate border and the prolonged expression of *pou5f3.2* and *sox3* in animal pole explants raised the possibility that loss of klf2 or klf17 was prolonging the pluripotency of blastula stem cells. We therefore asked if depletion of klf2 or klf17 would allow blastula explants to respond to neural inducing cues past the time when control explants can do so. Treatment with BMP inhibitor K02288 at stage 9 directs blastula explants to adopt a neural fate and express *sox2* or *nrp1* at stage 17 (Fig. 8 A,B). Depletion of either klf2 or klf17 does not impede the induction of either *sox2* (klf2MO: 97.8%, n=46; klf17MO: 100%, n=45; WT: 100%, *n*=43) or *nrp1* (klf2MO: 92.3%, *n*=39; klf17MO: 95.7%, n=47; WT: 100%, n=40) expression when BMP signaling is inhibited at stage 9 (Fig. 8 A,B). When BMP inhibition is initiated at stage 11.5, during gastrulation, cells are no longer competent to adopt a neural fate, as evidenced by a lack of *sox2* (97.1%, *n*=35) or *nrp1* (97.4%, *n*=38) expression. Strikingly, however, explants in which klf2 or klf17 have been depleted were able to adopt a neural fate when treated with K02288 at stage 9 as evidenced by expression of *sox2* (klf2MO: 92.9%, *n*=42; klf17MO: 95.1%, *n*=41) and *nrp1* (Klf2MO: 97.3%, *n*=37; Klf17MO: 97.2%, *n*=36) (Fig. 8 A,B).

**Fig 8.**
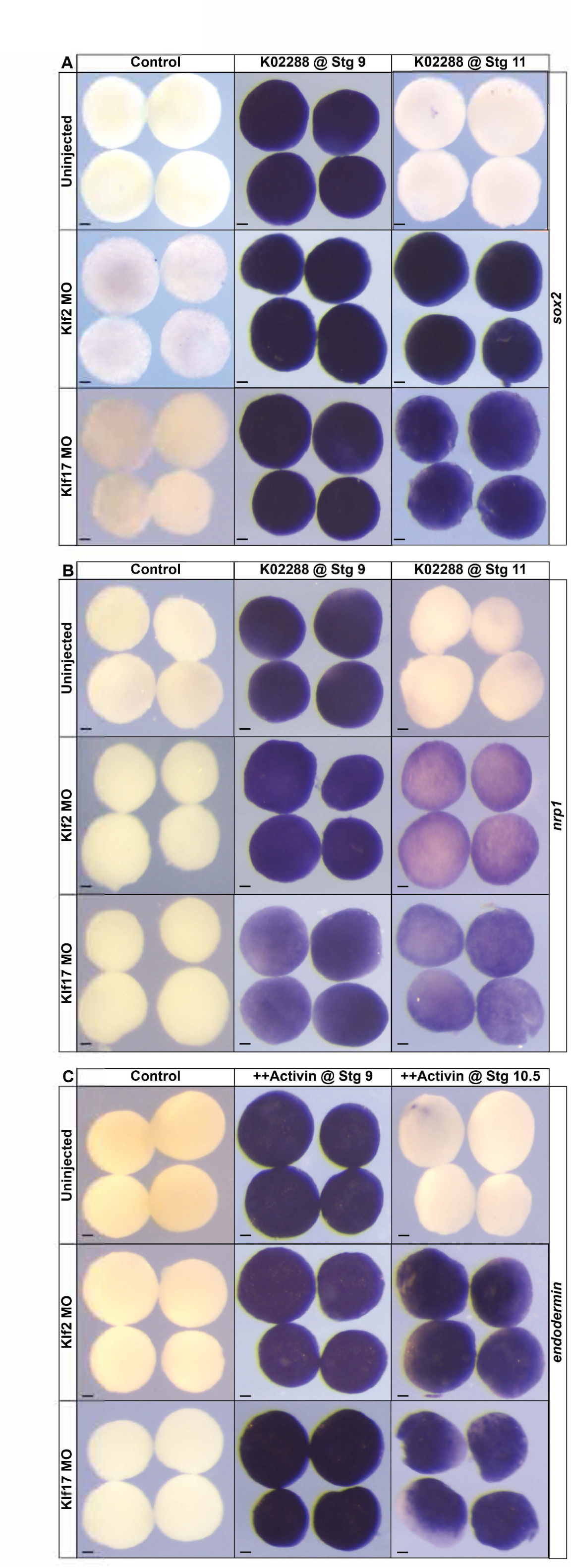
Knockdown of klf2 or klf17 extends the window of competency to adopt different states. (A-C) *In situ* hybridization examining the expression of neural markers (A) *sox2* and (B) *nrp1* in uninjected, *klf2* morphant, or *klf17* morphant explants treated with the BMP antagonist K02288 at either stage 9 or stage 11 and then collected for analysis at stage 17. (C) *In situ* hybridization of examining the expression of endodermal marker *endodermin* in uninjected, *klf2* morphant, or *klf17* morphant explants treated with high levels of activin at either stage 9 or stage 10.5 and then collected for analysis at stage 12. MO, morpholino; K02288, BMP inhibitor; CHIR, CHIR99021 (Wnt agonist); ++, high activin; scale bars: 100 µM.

To determine if depletion of klf2 or klf17 were prolonging pluripotency or only the competence to form neural progenitors, we next asked if loss of klf2 or klf17 could also prolonged the ability of these explants to form endoderm. Treatment of control explants or explants depleted for klf2 or klf17 with high levels of activin at stage 9 induced strong expression of the endoderm marker *endodermin at* stage 13 (klf2MO: 100%, *n*=26; klf17MO:100%, *n*=30; WT: 100%, *n*=36) (Fig. 8C). Treatment of control explants were with high activin at stage 10.5 failed to induce endoderm, however, indicating that cells has lost the competence to respond by this stage. By contrast, however, explants depleted for either klf2 or klf17 were still competent to for endoderm in response to high activin at stage 10.5, as evidenced by strong *endodermin* expression (klf2MO: 92.9%, *n*=28; klf17MO: 93.5%, *n*=31, WT: 0%, *n*=34) (Fig. 8C). These data further support a model whereby klf2 and klf17 function to restrict rather than promote pluripotency.

## Discussion

Recent work has shown that there is high degree of overlap between the GRNs of the neural crest and pluripotent blastula stem cells (Buitrago-Delgado et al. 2015; Zalc et al. 2021; Hovland et al. 2022; Schock, York, and LaBonne 2023). At the core of the pluripotency GRN circuitry are the factors identified by Yamanaka (pou5(Oct3/4), soxb1(sox2/3), klf4, Myc) as sufficient to reprogram somatic cells to a pluripotent state (Takahashi and Yamanaka 2006). These factors have been shown to be expressed in neural plate border and neural crest cells across multiple vertebrate species (Bellmeyer et al. 2003; Lavial 2007; Buitrago-Delgado et al. 2015; Lignell et al. 2017; Bhattacharya et al. 2018; Scerbo 2020; Zalc et al. 2021). Additionally, loss-of-function experiments have demonstrated that at least some of these key pluripotency factors are required for neural crest formation (Bellmeyer et al. 2003; Kurauchi, Izutsu, and Maeno 2010; Scerbo 2020; Zalc et al. 2021). Recent work in lamprey provided strong evidence that this shared pluripotency/neural crest GRN evolved at the base of the vertebrates (York et al. 2024).

A notable finding from that study was that, unlike mammals and other amniotes, lamprey *klf2/4* is not expressed in blastula stem cells or neural crest (York et al. 2024). Instead, it is *klf17* that is expressed in these stem cell populations. While *klf17* is also the most highly expressed klf factor in *Xenopus* neural crest, it is *klf2* that is the predominant *klf* factor expressed in pluripotent blastula cells. By contrast *klf4* is expressed in mouse neural crest cells (Pajanoja et al. 2023). Forced expression of klf4 has been shown to prevent ES cell differentiation (Zhang et al. 2010) similar to what we find for klf2 and klf17. While Zang et al., also proposed that Klf4 activity was required for both ES cell self-renewal and maintenance of pluripotency in mouse, more recent studies using inducible knockouts for *Klf2*, *Klf4*, and *Klf5* have shown that all three must be knocked out to completely block self-renewal (Yamane et al. 2018). Moreover, expression of any of the three can rescue pluripotency.

Interestingly, *KLF17* is expressed in the epiblast of pre-implantation human embryos (Blakeley et al. 2015) as well as in human naïve-like pluripotent stem cells (Takashima et al. 2014). As human pluripotent stem cells do not express *KLF2*, this would be consistent with a functional replacement by *KLF17* (Blakeley et al. 2015).

There is additional evidence, beyond control of pluripotency, for interchangeable functions of Klf factors during early development. In zebrafish both *klf2a/b* and *klf17* have been shown to regulate ectoderm and mesendoderm development (Gardiner et al. 2005; Kotkamp et al. 2014). In *Xenopus* inhibition of Klf4 function leads to failure of mesendoderm formation (Cao et al. 2012) and *Klf4* overexpression promotes neuroectoderm and endoderm formation. Interestingly, the expression patterns of *KLF2* and *KLF17* in human embryos are diametrically opposite to that of *klf2* and *klf17* in mouse (Yan et al. 2013; Blakeley et al. 2015). *klf17* is maternally provided in mouse eggs and its expression abolished around the eight-cell stage, whereas KLF17 becomes significantly upregulated in eight-cell human embryos, following zygotic genome activation (ZGA) (D’eng; Yan et al. 2013; Blakeley et al. 2015). Conversely, *klf2* is expressed from the two-cell stage, corresponding to mouse ZGA, and continues through to the blastocyst stage, whereas human *KLF2* is only expressed pre-ZGA (D’eng; Yan et al. 2013; Blakeley et al. 2015)

In the current study we find that loss of either klf2 or klf17 activity leads to expanded expression of both neural plate border and neural crest markers, although the increase is more pronounced for neural plate border. This suggests a role for these klf factors in controlling the extent to which the developing ectoderm retains pluripotency. Consistent with such a role, ectopic expression of either klf2 or kl17 inhibits expression of neural crest markers *snai2* and *foxd3*, whereas morpholino depletion of either factor significantly expands expression of *pou5f3.3* at neural plate stages.

We note that a previous study reported that loss of klf17 (then called neptune) led to loss rather than enhanced expression of neural crest markers (Kurauchi, Izutsu, and Maeno 2010). Those experiments are not directly comparable to the current study as their MOs were delivered throughout the embryo whereas here they were targeted to the ectoderm. Also, we have found that partial depletion of *klf17* leads to significantly increased expression of endogenous *klf17* (unpublished data), so it is possible that study may have inadvertently been generating a gain-of-function phenotype.

Consistent with a role in inhibiting pluripotency, we found that forced expression of klf2 or klf17 prevented blastula explants from transiting to an epidermal, mesodermal or endodermal state. By contrast depletion of either klf2 or klf17 in these explants prolongs the expression of pluripotency factors *sox3* and *pou5f3.2*. A consequence of this is decreased expression of *krt12.4* at stage 13 (Fig.7C). Strikingly, however it does not affect expression of the neural plate border factor *pax3*, emphasizing the connectedness of the pluripotency and neural plate border/neural crest GRNs. Given the extended *sox3* and *pou5f3.2* expression we asked if klf depletion would prolong functional pluripotency and found that the time window during which both the neural and endodermal states could be induced was indeed extended.

Our results suggest that klf2 and klf17 are the primary klf factors that regulate pluripotent blastula and neural crest stem cells in *Xenopus*, whereas *klf4* is not significantly expressed in these populations. By contrast in amniotes such as mouse and human, klf4, one of the classical Yamanaka factors, is a key regulator of pluripotency in ES cells, although klf2 and klf17 also play roles. The pluripotency promoting activities of klf factors in amniotes contrasts with our findings that in *Xenopus,* klf2 and klf17 restrain pluripotency. Given this divergence, we wished to gain insights into what the ancestral role of these factors might have been in stem vertebrates. Accordingly, we turned to lampreys, one of two extant jawless vertebrates.

Shared features of lamprey and jawed vertebrates likely represent features of their last common ancestor. Lamprey possess a *klf17* and a *klf2/4* factor however only *klf17* is expressed in blastula animal pole cells and neural crest (York et al. 2024). Given these conserved phenotypes, we sought to trace the stepwise evolutionary origins of klf17 activity in vertebrates by performing molecular phylogenetic analysis and chromosomal synteny comparisons (Fig. S4). Consistent with recent whole-genome analyses in hagfish, lamprey, and other vertebrates, our results suggest that a whole-genome duplication event gave rise to klf17 and klf2/4 in ancestral cyclostomes, features that are in extant lampreys. Importantly, synteny comparisons between *Xenopus* and lamprey show that both *klf2* and *klf4* in *Xenopus* can be mapped to the single lamprey *klf2/4* ortholog (Fig. S3), suggesting that *klf2* and *klf4* arose by tandem gene duplication in stem gnathostomes. These results suggest that the neural crest/pluripotency restraining activities of klf17 transcription factors were present in the last common ancestor of jawed and jawless vertebrates and that they evolved new functions in the lineage leading to extant amniotes. They also support a model in which *klf2*, *klf4*, and *klf17* emerged in jawed vertebrates through stepwise evolution via gene duplication of an ancestral *klf2/4/17* gene present in invertebrate chordates.

## MATERIALS AND METHODS

### Embryological Methods

Wild type *Xenopus laevis* embryos were staged and collected in accordance with standard methods (Zahn et al. 2022). *In situ* hybridizations were performed on embryos and explanted animal caps using previously described methods (LaBonne 1998). Microinjection of mRNA (Ambion, mMessage mMachine SP6 Transcription Kit) or morpholino (Gene Tools) was done in 1-4 cells at the 2-8 cell stage as previously described (Lee et al. 2012). Approximately 10-25 ng of translation-blocking morpholinos (Gene Tools) were injected per cell. Manipulated embryos were then cultured in 0.1x Marc’s Modified Ringer’s Solution (MMR) [0.1 M NaCl, 2 mM KCl, 1 mM MgSO_4_, 2 mM CaCl_2_, 5 mM HEPES (pH 7.8), 0.1 mM EDTA] until being collected or dissected for animal cap explant assays. All animal cap explants were manually dissected during the early blastula stage and then cultured in 1xMMR until collection. For activin experiments, animal cap explants were dissected and immediately cultured in 1xMMR with 0.1% bovine serum albumin (BSA) and recombinant activin protein (R&D systems) at a final concentration of 20-40 ng/mL for mesoderm induction and 100 ng/mL for endoderm induction. For neural induction of animal cap explants, BMP signaling activity was inhibited with the small molecule inhibitor K02288 (Sigma) at a final concentration of 20 μM in 1xMMR as previously described (Johnson et al. 2022). Manipulated embryos and/or explants were fixed in 1x MEM [100mM MOPS (pH 7.4), 2mM EDTA, 1mM MgSO_4_] with 4% formaldehyde and dehydrated in methanol prior to *in situ* hybridizations. Results shown are representative of a minimum of three biological replicates.

#### DNA constructs

Full length *Xenopus* Klf2 and Klf17 were obtained from the Xenopus ORFeome (www.xenbase.org/reagents/static/orfeome.jsp) and subcloned into pCS2 vectors for synthesis of mRNA for microinjections. For gain of function experiments, each respective mRNA was injected together with mRNA encoding the lineage tracer β-gal. The 3’UTR regions of *klf2* and *klf17* were amplified from genomic DNA isolated from WT embryos and subcloned into pGEM-T vector for synthesis of RNA probes. The morpholino antisense oligonucleotides against the 5′UTR coding regions of Xenopus klf2 (5’-GAGAATGGTCTCGCTCAGAGCCATC) or klf17 (5’-GGGTTGAGAAAGCCACACTCATCCT) conjugated to FITC were validated by co-injecting it with an epitope tagged versions of their respective mRNA for western blot analysis.

### Western Blot

Five whole embryos were lysed in 1% NP-40 supplemented with protease inhibitors (Complete Mini, EDTA-free tablet (Roche), Leupeptin (Roche), Aprotinin (Sigma), and phenylmethylsulfonyl fluoride (PMSF; Sigma)). SDS page and western blot were used to detect proteins. The following antibodies were used: c-Myc 9E10 (1:3000; Santa Cruz; sc-40); actin (1:5000; Sigma; A2066); IRDyes (1:20,000 mouse-800 CW; rabbit-680 TL) and the Odyssey platform (LI-COR Biosciences) were used to detect proteins. Results are representative of a minimum of three biological replicates.

#### Hybridized Chain Reaction

Hybridized chain reaction (HCR) methodologies are adapted from (Choi et al., 2018). Whole embryos were hybridized with DNA probe sets for *klf17*, *pax3*, *snai2*, *foxd3*, and *sox3* (Molecular Instruments) and incubated overnight at 37 °C. Probe was removed, samples washed, and then incubated overnight with DNA hairpins labeled with Alexa488, Alexa 647 or Alexa 546 (Molecular Instruments). Unbound hairpins were removed via 5x SSCT washes followed by PBS washes and incubated in DAPI (1:5000; Life Technologies). Samples were mounted and imaged using a Nikon C2 upright confocal with two GaAsP detectors and four standard laser lines with either 4x or 10x objectives.

#### Line Profile Analysis

Line profile measurements of *klf17, pax3 and snai2* were made on multi-channel maximum intensity projections from confocal files using the python packages scikit-image(Van Der Walt et al. 2014). Intensity profiles for each fluorescent channel were measured using the profile_line function from scikit-image along a user defined line with endpoints chosen manually using the python package, mpl_point_clicker (https://github.com/ianhi/mpl-point-clicker). The intensity profile was then normalized to a range between 0 and 1 by subtracting the minimum intensity value and dividing by the range (maximum - minimum intensity) for each channel. In order to compare multiple line profiles from each image the average *pax3* or *snai2* intensities were first fit to an exponential curve and the peaks of each centered at zero. After curve fitting the common x-values (pixel distance) were identified by calculating the intersection of the x-values across all measurement curves and trimmed to only include common x-values. The mean and standard error of the normalized intensities of *klf17* in combination with either *pax3* or *snai2* were then calculated and plotted across the x-values.

### Animals

All animal procedures were approved by the Institutional Animal Care and Use Committee, Northwestern University, and are in accordance with the National Institutes of Health’s Guide for the Care and Use of Laboratory Animals.

## Acknowledgements

We thank S. Miehls and the staff of the Hammond Bay Biological Station for shipment of lampreys and members of the lab for helpful discussions. The authors acknowledge the contributions of Xenbase (https://www.xenbase.org/xenbase) and the National Xenopus Resource (https://www.mbl.edu/research/resources-research-facilities/national-xenopus-resource).

## Funding

Funding for the study was received from Life Sciences Research Foundation postdoctoral fellowship (J.R.Y.), National Institutes of Health (grant nos. R01GM116538 (C.L.), National Science Foundation (grant no. 1764421 to C.L.) and Simons Foundation (grant no. SFARI 597491-RWC to C.L.).

## Notes

### Competing Interest Statement

The authors have declared no competing interest.

