## Supplemental data for "Krüppel-like Factors Play Essential Roles in Regulating Pluripotency and the Formation of Neural Crest Stem Cells"

**Supplemental Figure. 1:**

**A**

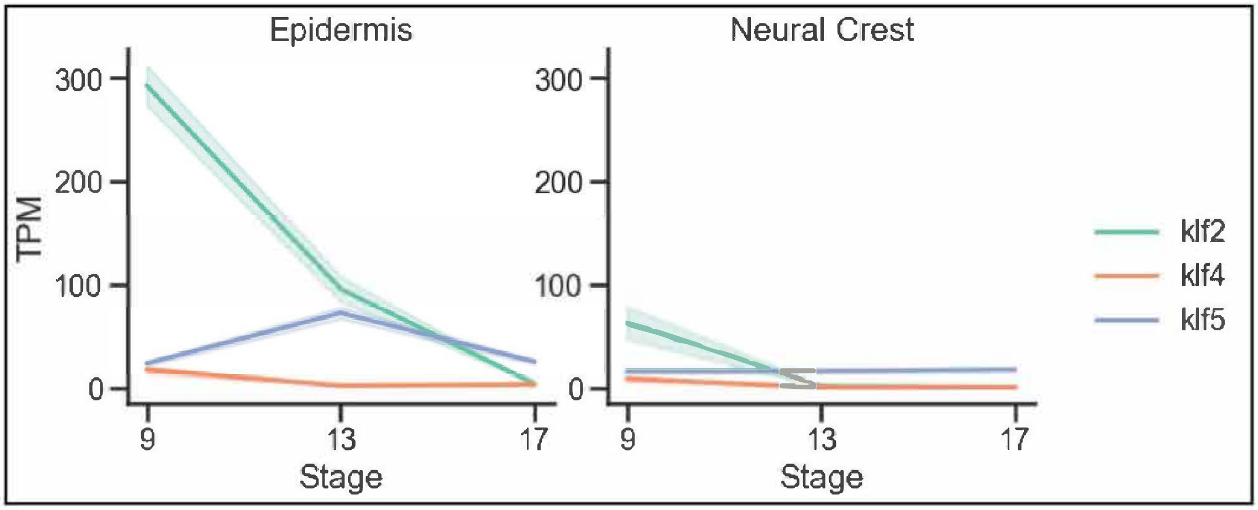


**B**

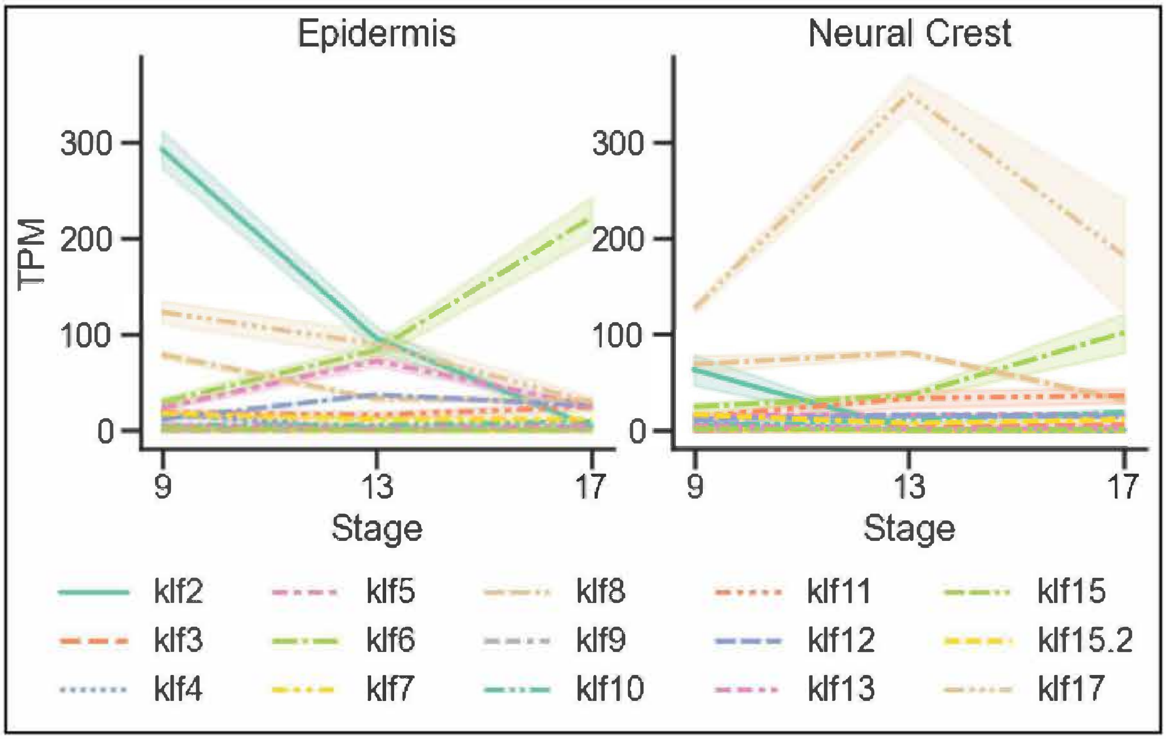


**Sup. Fig. 1: TPM plots of klf factors in epidermal and neural crest explants.**
(A-B) Plots of the average TPMs of Klf factors in epidermis and neural crest explants at blastula (stage 9), early neurula (stage 13), and late neurula (stage 17) stages. Shading represents the standard error of mean. (A) Plots of *klf2, klf4* and *klf5* TPMs. (B) Plots of all 15 klfs in the *Xenopus laevis* genome.

### Supplemental Figure 2:

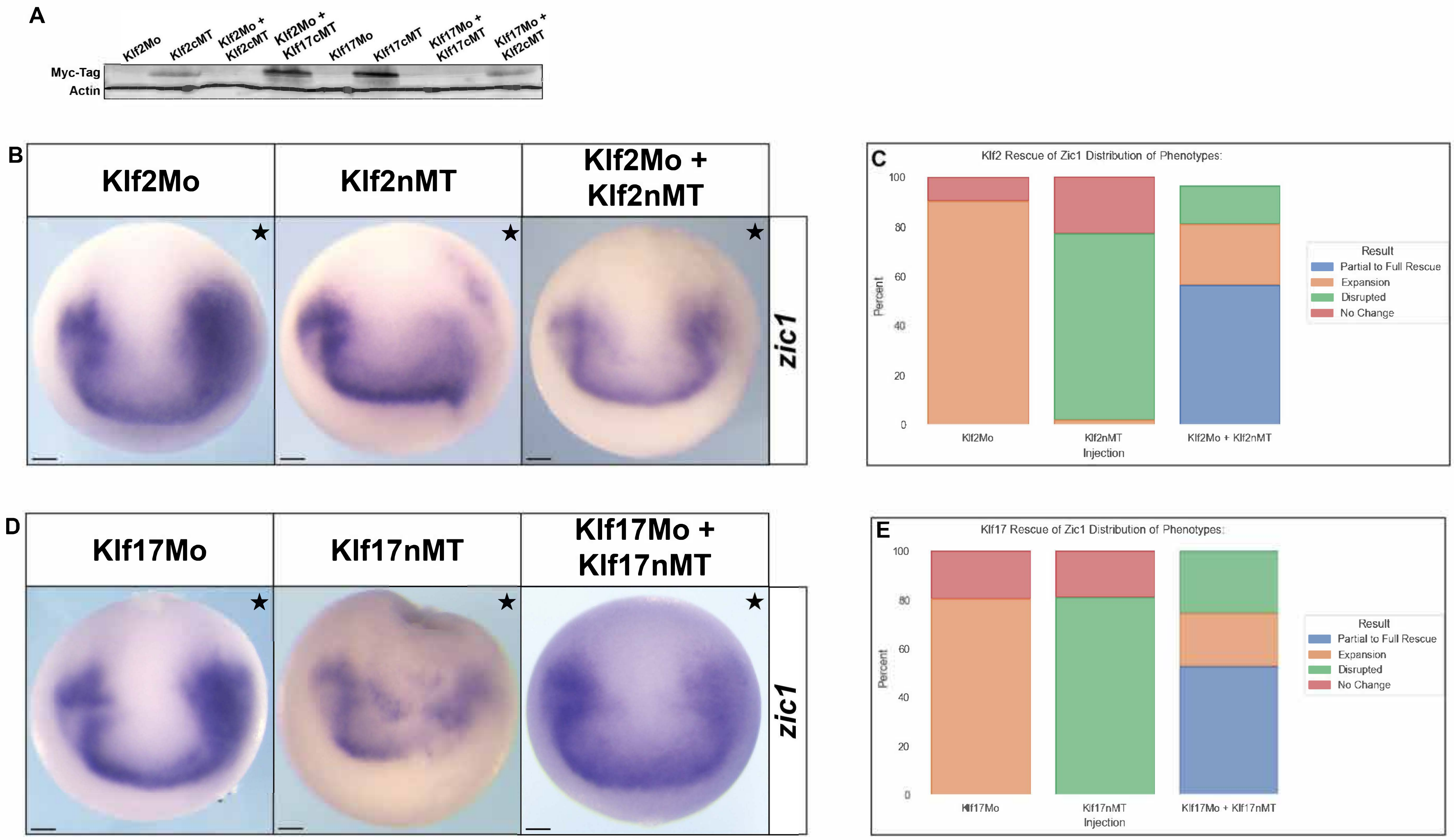


**Sup. Fig. 2: Klf2 and Klf17 Morpholino Validation.** (A) Western blot validation of Klf2 and Klf17 morpholinos. (B) *In situ* hybridizations showing the rescue of *zic1* expression in *klf2*-morphant embryos co-injected with epitope tagged *klf2* mRNA (C) Stacked bar graphs of the percent distribution of phenotypes for the *klf2* rescue of *zic1* expression (Klf2Mo n=52; klf2-nMT(n=92; Klf2Mo + klf2-nMT n= 90). (D) *In situ* hybridizations showing the rescue of *zic1* in *klf17*-morphant embryos co-injected with epitope tagged *klf17* mRNA. (E) Stacked bar graphs of the percent distribution of phenotypes for the *klf17* rescue of *zic1* expression (Klf17Mo n=62; klf17-nMT n=63; Klf17Mo + klf17-nMT n=55) . cMT, C-terminal myc tag; nMT, N-terminal myc tag; MO, morpholino.

### Supplemental Figure 3:

A

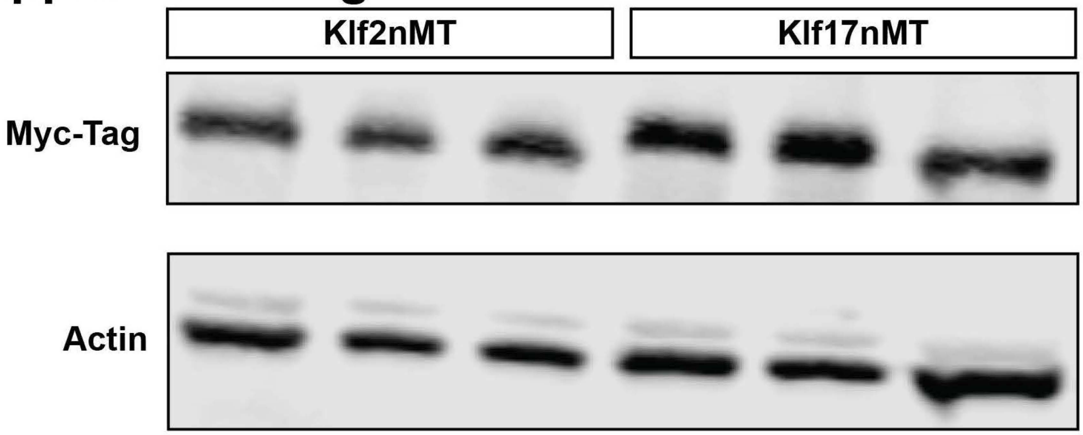


**Sup. Fig. 3: Western Blot to Validate Protein Levels.** (A) Western blot of three replicates of epitope tagged *klf2* and *klf17* used in gain of function experiments at consistent levels. nMT, N-terminal myc tag

### Supplemental Figure 4:

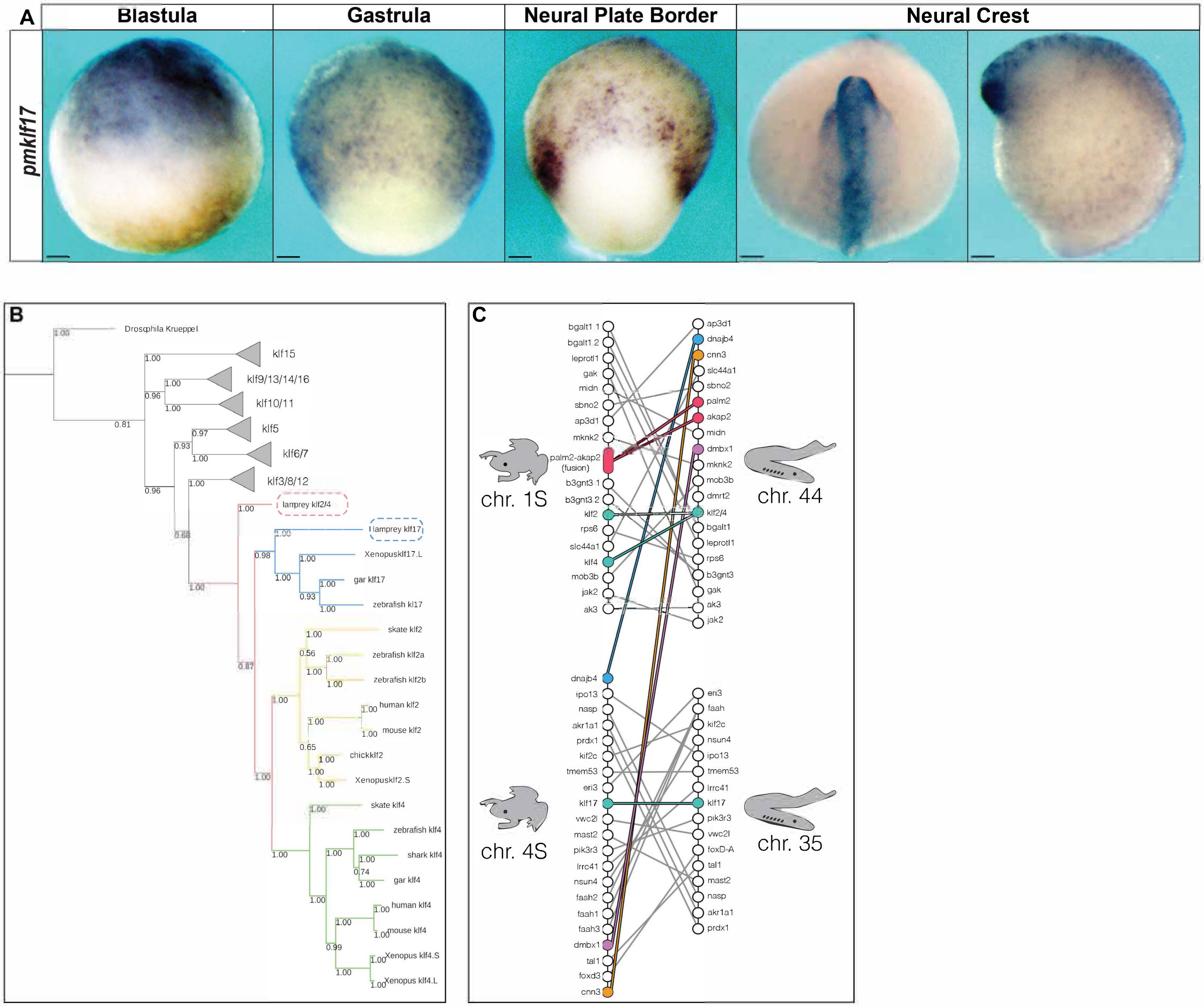


**Sup. Fig. 4: Evolution and Conservation of Klf factors.** (A) *In situ* hybridizations of *klf17* expression in the blastula, gastrula, neural plate border and neural crest of wild type sea lamprey embryos. (B) Phylogenetic tree of Klf factors across several species. (C) Synteny analysis of *klf2* and *klf17* in *Petromyzon marinus* (sea lamprey) and *Xenopus laevis*. pm, *Petromyzon marinus.*
